# Engineering SARS-CoV-2 cocktail antibodies into a bispecific format improves neutralizing potency and breadth

**DOI:** 10.1101/2022.02.01.478504

**Authors:** Zhiqiang Ku, Xuping Xie, Jianqing Lin, Peng Gao, Abbas El Sahili, Hang Su, Yang Liu, Xiaohua Ye, Xin Li, Xuejun Fan, Boon Chong Goh, Wei Xiong, Hannah Boyd, Antonio E. Muruato, Hui Deng, Hongjie Xia, Zou Jing, Birte K. Kalveram, Vineet D. Menachery, Ningyan Zhang, Julien Lescar, Pei-Yong Shi, Zhiqiang An

**Affiliations:** Texas Therapeutics Institute, Brown Foundation Institute of Molecular Medicine, University of Texas Health Science Center at Houston, Houston, TX 77030, USA; Department of Biochemistry and Molecular Biology, Institute for Human Infection and Immunity, Sealy Institute for Vaccine Sciences, Sealy Center for Structural Biology & Molecular Biophysics, Department of Pharmacology & Toxicology, University of Texas Medical Branch, Galveston, TX 77555, USA; NTU Institute of Structural Biology and School of Biological Sciences, Nanyang Technological University, 636921, Singapore; Antimicrobial Resistance Interdisciplinary Research Group, Singapore-MIT Alliance for Research and Technology Centre, 138602, Singapore; Department of Microbiology & Immunology, University of Texas Medical Branch, Galveston, TX, 77555, USA

## Abstract

One major limitation of neutralizing antibody-based COVID-19 therapy is the requirement of costly cocktails to reduce antibody resistance. We engineered two bispecific antibodies (bsAbs) using distinct designs and compared them with parental antibodies and their cocktail. Single molecules of both bsAbs block the two epitopes targeted by parental antibodies on the receptor-binding domain (RBD). However, bsAb with the IgG-(scFv)_2_ design (14-H-06) but not the CrossMAb design (14-crs-06) increases antigen-binding and virus-neutralizing activities and spectrum against multiple SARS-CoV-2 variants including the Omicron, than the cocktail. X-ray crystallography and computational simulations reveal distinct neutralizing mechanisms for individual cocktail antibodies and suggest higher inter-spike crosslinking potentials by 14-H-06 than 14-crs-06. In mouse models of infections by SARS-CoV-2 and the Beta, Gamma, and Delta variants, 14-H-06 exhibits higher or equivalent therapeutic efficacy than the cocktail. Rationally engineered bsAbs represent a cost-effective alternative to antibody cocktails and a promising strategy to improve potency and breadth.

## Introduction

The COVID-19 pandemic has ravaged the world with unprecedented health, social and economic losses^1^. Vaccination is among the most effective countermeasures but not sufficient to end the pandemic due to challenges such as limited global access, vaccine hesitancy, and waning effectiveness against variants^2–4^. Effective treatments are necessary for the patients, unvaccinated populations and immunocompromised people who cannot generate protective immunity after vaccination^5^.

Neutralizing antibodies have proved to be effective against COVID-19. The RBD of SARS-CoV-2 spike protein (S) directly contacts the cellular receptor angiotensin-converting enzyme 2 (ACE2). It is the target of the most potent neutralizing antibodies^6^. However, drug resistance rapidly arises with antibody monotherapies regardless of neutralizing potency and epitope conservation of the antibodies^7^. Emerging SARS-CoV-2 variants of concern (VOC), such as the Beta and Gamma, have evolved RBD mutations that escape from neutralization by many single antibodies and some combined antibodies with overlapping epitopes^8^. Rationally designed antibody cocktails, which cover non-overlapping epitopes, can reduce SARS-CoV-2 escape mutations and expand neutralizing coverage of emerging variants^9,10^. Three antibody cocktails have received approval for emergency use, and several candidates are in advanced stages of clinical trials. Despite the encouraging progress, antibody cocktail approaches increase manufacturing costs and require high dose infusion in patients^11^, making it challenging to have a global impact on pandemic response^12^. Recently, a wide range of antibodies has dramatically or completely lost neutralization against the Omicron variant^13–16^, making the FDA to limit the use of two approved antibody cocktails.

Bi-specific antibodies (bsAbs) are an emerging drug modality designed to combine the binding specificities of two antibodies into one molecule. With different designs, bsAbs can be engineered into diverse formats with varied valencies. One attractive feature for bsAbs is their potential to display novel functionalities that do not exist in mixtures of parental antibodies^17^. For example, engineered HIV-1 neutralizing bsAbs in the CrossMAb format have enhanced virus-neutralizing potency and breadth compared with the mixtures of parental antibodies^18,19^. With the same CrossMAb design, a SARS-CoV-2 bsAb (CoV-X2) exhibits a neutralizing activity superior to one parental antibody and similar to the other parental antibody^20^, suggesting the need to test other bsAb designs for improvement of bsAb functions. *In vitro* and *in vivo* comparisons of bsAbs with parental antibodies and the cocktail, which are lacking in previous studies, will provide more insights for developing efficacious bsAb-based COVID-19 therapeutics.

We have previously identified two SARS-CoV-2 neutralizing antibodies, called CoV2-06 and CoV2-14, respectively, recognizing non-overlapping RBD epitopes and preventing escape mutations as a cocktail^21^. In this study, we engineered the two antibodies into two bsAbs, one using the CrossMAb design and the other using the IgG-(scFv)_2_ design. Using biochemical, structural, and virological assays, we demonstrate that the IgG-(scFv)_2_ design, but not the CrossMAb design, enhances neutralizing potency and spectrum against multiple SARS-CoV-2 variants in comparison with parental antibodies and the cocktail.

## Results

### Engineering of bispecific antibodies

We sought to construct bsAbs to combine the utility of CoV2-06 and CoV2-14 into one single molecule. To explore whether and how the design of formats affect the functions of bsAbs, we engineered two bsAbs with distinct features: one bsAb (14-H-06) is in the tetravalent format using the IgG-(scFv)_2_ design, and the other bsAb (14-crs-06) is in the bivalent format using the CrossMAb design (**Fig. 1a**). The two bsAbs were produced by transient expressions in Expi293F cells with high yields (>500 μg/ml). After a single-step Protein A chromatography purification, the bsAbs were showed >95% purities and correctly assembled as analyzed by size-exclusion chromatography (SEC) (**Fig. 1b**). To test whether the bsAbs block the two epitopes targeted by CoV2-06 and CoV2-14, we performed an in-tandem Bio-Layer Interferometry (BLI) based binding assay (**Fig. 1c**). Both 14-H-06 and 14-crs-06 bound to RBD and blocked the subsequent binding of CoV2-06 and CoV2-14 (**Fig. 1d**). In contrast, pre-binding of RBD by CoV2-06 or CoV2-14 did not abolish subsequent binding of 14-H-06 or 14-crs-06 (**Fig. 1e**). These results indicate that the bsAbs are successfully engineered and both of them block the two RBD epitopes simultaneously as single molecules.

**Fig. 1.**
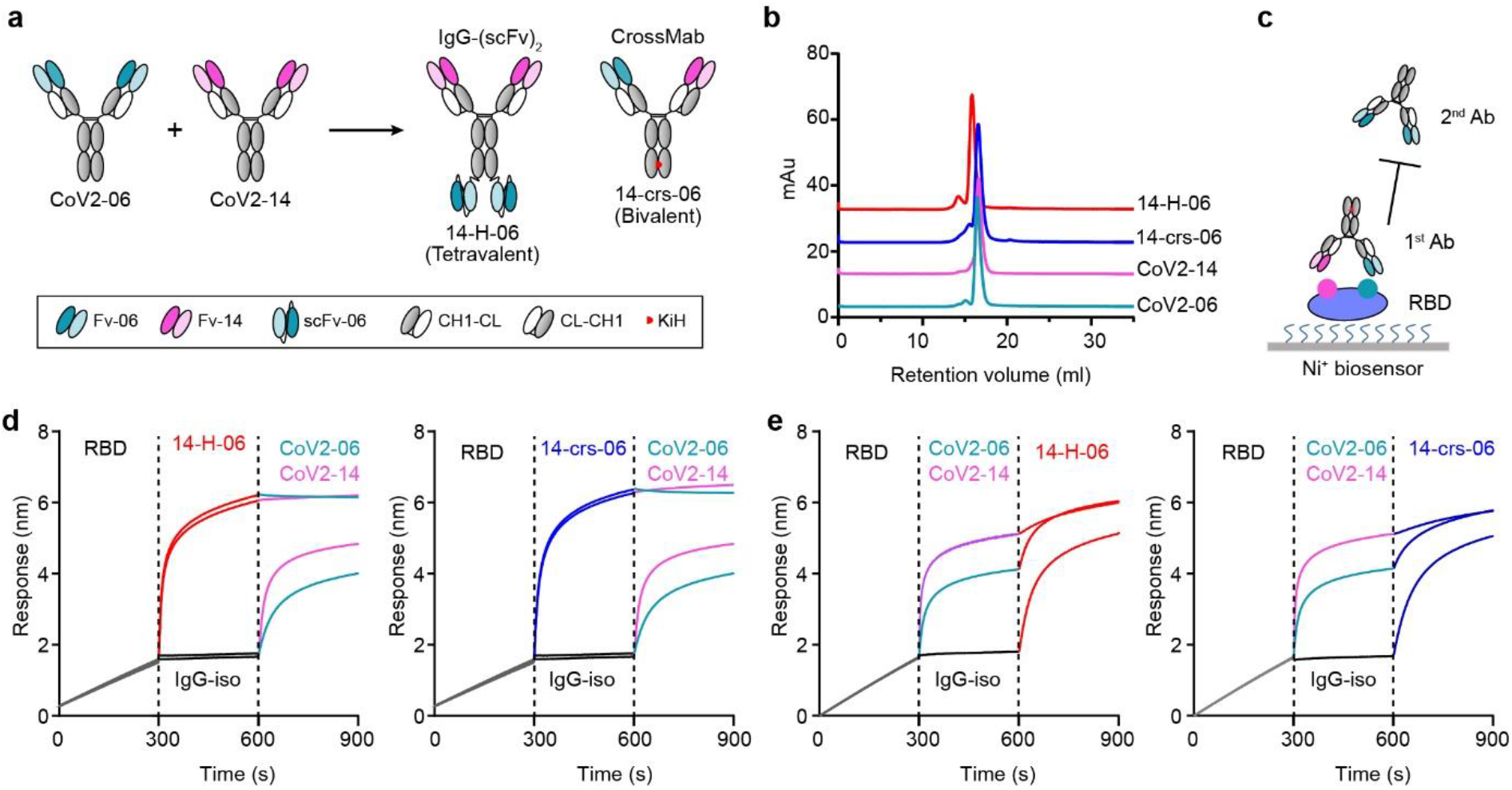
Engineering of bispecific antibodies. **a**, Schematic diagram of engineering bispecific antibodies. A tetravalent bsAb (14-H-06) and a bivalent bsAb (14-crs-06) were engineered from two parental IgGs that bind to two distinct epitopes on the RBD. The modules for antibody engineering are illustrated in the box. Fv: variable fragment; scFv: single-chain Fv; CH1-CL: constant region 1 for heavy chain (CH1) and constant region for light chain (CL); CL-CH1: the crossover format of CH1-CL; KiH: the Knob-into-Hole design with the S354C+T366W mutations (Knob) in the heavy chain CH3 region on one arm and the Y349C+T366S+L368A+Y407V mutations (Hole) in the CH3 region on the other arm. **b**, Purities of indicated antibodies were analyzed by SEC. **c**, The in-tandem BLI-based assay to test antibody blocking of RBD epitopes. The His-tagged RBD was captured onto the Ni-NTAbiosensors. The first antibodies, either bsAbs or parental antibodies, were used to bind the RBD. The second antibodies were tested for their abilities to bind pre-blocked RBD. **d-e**, The epitope pre-blocking effects using bispecific antibodies 14-H-06 (left) and 14-crs-06 (right) as 1^st^ antibodies and parental antibodies as 2^nd^ antibodies (**d**), or using parental antibodies as 1^st^ antibodies and the bispecific antibodies 14-H-06 (left) and 14-crs-06 (right) as 2^nd^ antibodies (**e**). The segments of each sensorgram were color coded to show individual binding events.

### Enhanced antigen binding and virus neutralization for 14-H-06

We characterized the antigen-binding properties of the two bsAbs and the two parental antibodies using BLI-based kinetic assays. To measure the affinity binding, we immobilized antibodies onto protein A biosensors and used soluble His-tagged RBD (RBD-His) as the analyte. Both bsAbs bound to RBD with affinity K_D_ values in the low nanomolar range and comparable to the two parental antibodies (**Extended Data Fig. 1a-b**). The result is consistent with the intrinsic binding strength of the one-to-one interaction for the Fab, or scFv, to the RBD. To measure the avidity binding, we immobilized Ni-NTA biosensors with RBD-His at different concentrations and used antibodies as the analyte (**Fig. 2a and Extended Data Fig. 1c-e**). Avidity represents the combined strength of all binding sites on an antibody molecule and often manifests as decrease of dissociation from tethered antigens^22^. Accordingly, as the concentration of RBD for immobilization increased from 40 ng/ml to 1000 ng/ml, the tetravalent antibody14-H-06 showed a greater increase of avidity binding (K_D_ values changed from 1.35 nM to <0.001 nM, over 1350-fold) than the bivalent antibodies 14-crs-06 (8.6-fold change), CoV2-06 (46-fold change) and CoV2-14 (2.0-fold change) (**Fig. 2b**). The increased avidity binding was due to much slower dissociation of 14-H-06 from the RBD than other antibodies, which were demonstrated by its larger fold changes of the 1/K_dis_ values (**Fig. 2c**). These results indicate that 14-H-06 enhances antigen binding activity with stronger avidity effects than the 14-crs-06.

**Fig. 2.**
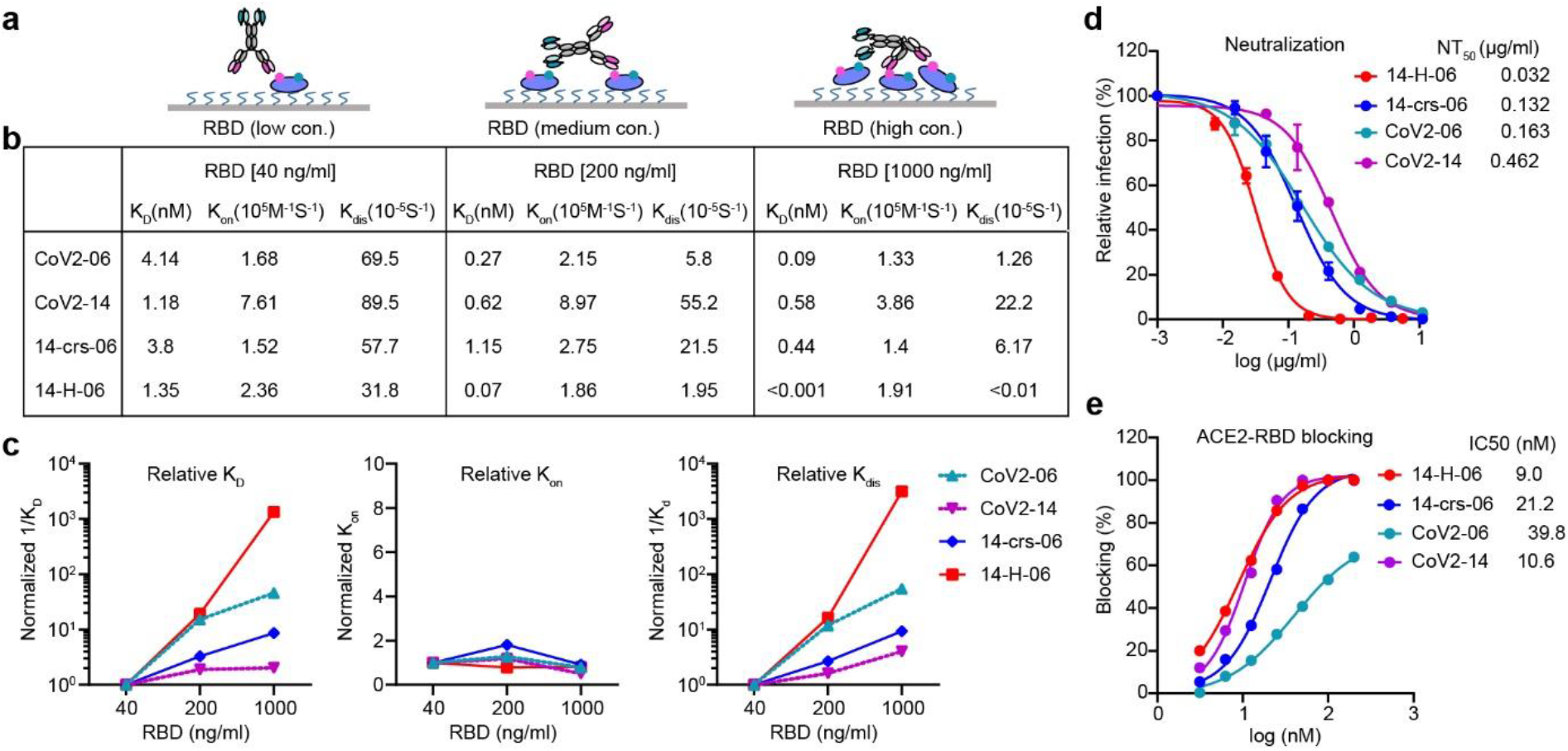
Enhanced avidity binding and virus neutralization for 14-H-06. **a**, A diagram showing the binding models of 14-H-06 to tethered RBD antigen at low (left), medium (middle) and high (right) concentrations. The avidity effects manifest as multivalent interactions between an antibody and multiple RBDs. **b**, Summary of the association (K_on_), dissociation (K_dis_) and avidity (K_D_) of indicated antibodies at indicated concentrations RBD. **c**, The plots of relative association, dissociation and avidity for each antibodies. The relative values for each antibodies were obtained by normalizing the values of 1/K_D_, K_on_ and 1/K_dis_ at RBD concentrations of 200 ng/ml and 1000 ng/ml against the corresponding values at RBD concentration of 40 ng/ml. **d**, Neutralization titrations of indicated antibodies against live SARS-CoV-2 on Vero E6 cell. Data points are from duplicate wells. **e**, Antibody blocking of RBD interaction with ACE2 determined by the BLI assay.

We compared the neutralizing activities of the bsAbs and parental antibodies using authentic SARS-CoV-2 virus^21^. Antibody 14-crs-06 neutralized SARS-CoV-2 with a half-maximal neutralizing titer (NT_50_) of 0.132 μg/ml, which was similar to CoV2-06 (NT_50_=0.163 μg/ml) and 3.5-fold better than CoV2-14 (NT_50_=0.462 μg/ml). This result is consistent with the trend that observed in CoV-X2, a previously reported SARS-CoV-2 bsAb with the same CrossMAb design^20^. In contrast, 14-H-06 neutralized SARS-CoV-2 with an NT_50_ of 0.032 μg/ml, which was 5.1-fold and 14.4-fold more potent than CoV2-06 and CoV2-14, respectively (**Fig. 2d**). To understand whether the two bsAbs alter the blocking activity against RBD binding to ACE2, we performed a BLI-based competition assay^23^. Antibody 14-H-06 blocked the RBD and ACE2 interaction with a half-maximal inhibition concentration (IC_50_) of 9.0 nM, which is similar to the IC_50_ of CoV2-14 (10.6 nM) and slightly lower than the IC_50s_ of 14-crs-06 (21.2 nM) and CoV2-06 (39.8 nM) (**Fig. 2e**). These results indicate that the avidity binding, but not the steric hindrance with ACE2, contributes to the improvement of neutralizing activity for 14-H-06 over 14-crs-06 and parental antibodies.

### Inter-spike crosslinking potential for 14-H-06

The avidity binding of an antibody offers the opportunity to engage multiple spikes through crosslinking, which is an extra line of neutralizing mechanisms for certain RBD-targeting antibodies^24^. We sought to compare the potentials for inter-spike crosslinking by the bsAbs and parental antibodies. In a BLI-based sandwich assay, RBD-His was immobilized onto Ni-NTA biosensors to capture antibodies, followed by incubation with Fc-tagged RBD (RBD-Fc). After RBD-His capturing, 14-H-06 showed much stronger binding to RBD-Fc than did 14-crs-06 and the two parental antibodies (**Extended Data Fig. 2**). The result indicates that 14-H-06 can engage more RBDs simultaneously than 14-crs-06 and parental antibodies through the four binding moieties, suggesting a higher potential for inter-spike crosslinking.

The orientations of Fab binding to RBD affect an IgG’s potential for inter-spike crosslinking^24^. We used X-ray crystallography and determined the complex structure of the Fab of CoV2-06 (Fab06) and RBD at a resolution of 2.89 Å (**Table S1**). The atomic details of interactions established at the binding interface between Fab06 and RBD showed that VH residues N32, W34 from CDR-H1, S55 from CDR-H2, and T104 from CDR-H3 interact with RBD residues N450, K444, Y449 and R346 while the VL residues N33 from CDR-L1 and D52 from CDR-L2 interact with RBD residues T345 and R346, respectively (**Fig. 3a**). The interactions revealed by X-ray crystallography are fully consistent with epitope mapping results reported in our previous study^9^. Next, we used the Fab06/RBD crystal structure (this work) to perform a superposition with two cryo-EM structures of the spike trimer: one where the three RBD adopt the down conformation and the other with one RBD up and two RBD down. As no steric hindrance was observed, this superposition suggests that Fab06 could bind to RBDs regardless of their down/up conformation in the spike trimer (**Fig. 3b, left**), which supports the ability of inter-spike crosslinking for bispecific antibodies incorporating the CoV2-06 paratope^24^. Although we were able to obtain the structure of the free Fab14 (**Extended Data Table 1**), so far, our attempts to use X-ray diffraction to determine a Fab14/RBD crystal structure have not been successful and work with Cryo-EM to determine this structure is in progress. Meanwhile, we used the X-ray structures of Fab14 and RBD for docking using Haddock 2.4 guided by previous epitope mapping results^9^. Docking suggests that Fab14 can only bind RBD in the up confirmation as, in the down conformation, Fab14 would clash with an adjacent RBD domain (**Fig. 3b, right**). Interestingly, Fab06 has little while Fab14 has large steric clash with ACE2 (**Fig.3c**). The binding epitopes and orientations indicate that the major neutralizing mechanisms for individual cocktail antibodies are different: CoV2-06 through crosslinking of spikes and CoV2-14 through ACE2 blocking. We used the molecular dynamics (MD) method to model the structures of bsAbs and superposed them with RBDs in the spike. The result shows that both bsAbs could simultaneously engage multiple RBDs in different spike trimer (**Fig. 3d**). However, the maximum number of spikes can be crosslinked by the two bsAbs were different when binding to RBDs adopting various combinations of up and down conformations. As summarized in **Fig. 3e**, the tetravalent 14-H-06 can crosslink more spikes than the bivalent 14-crs-06 and parental antibodies in all possible scenarios.

**Fig. 3.**
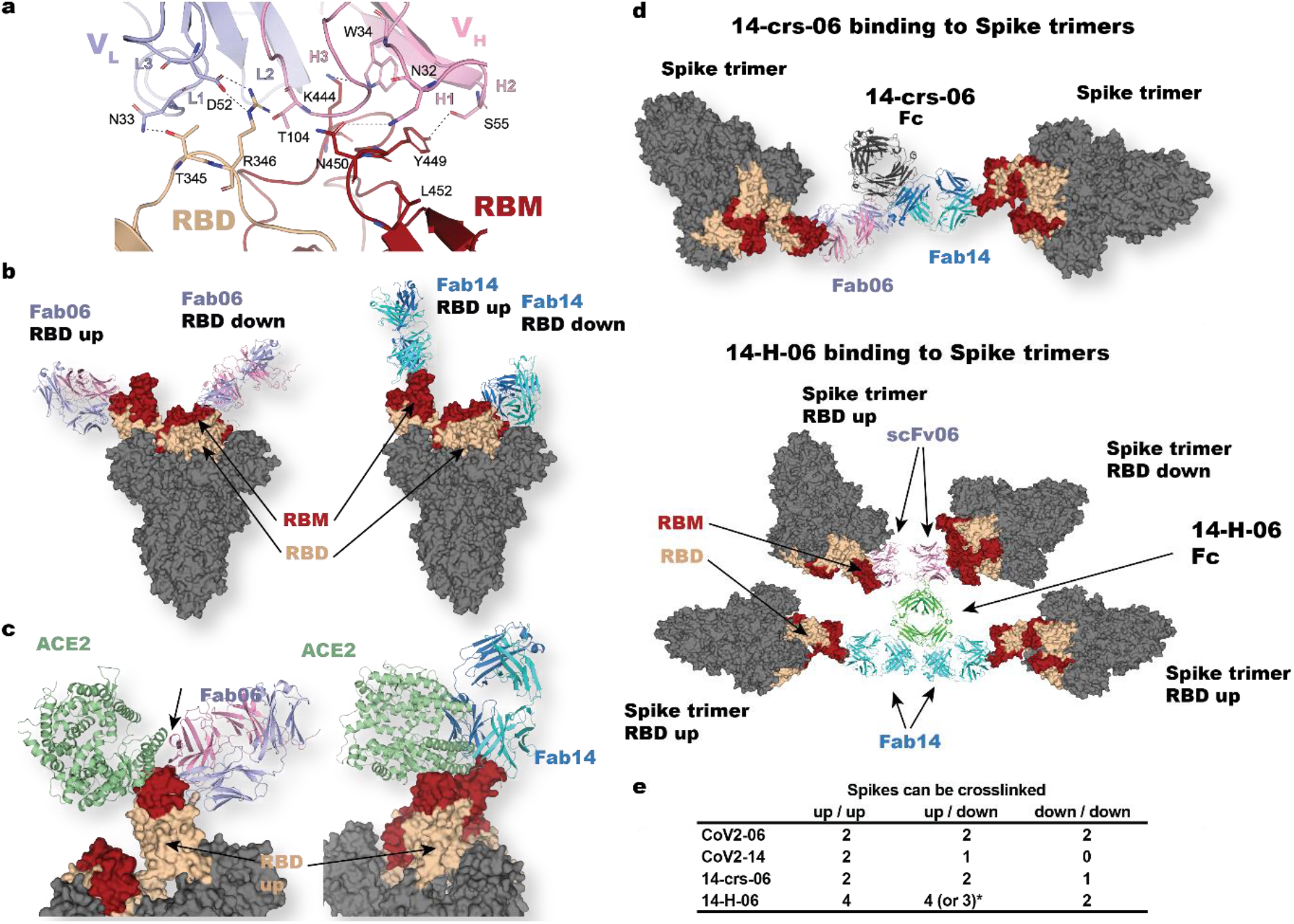
Binding and inter-spike crosslinking potential for 14-H-06. **a**. Atomic details at the binding interface between Fab-06 and RBD as revealed by X-ray crystallography. The VH and VL of Fab-06 are colored in red and blue, respectively and RBD is colored in yellow. Dashed lines indicate polar interactions between Fab-06 and RBD. **b**. Binding models for Fab-06 (left) or Fab-14 to RBDs (either in up or down conformation) in the context of the complete spike trimer. **c**. Based on the epitope they recognize, both Fab-06 and Fab-14 introduce steric hindrance (indicated by arrows) with ACE2 binding to RBD in the up conformation. **d**, Cross-links induced by the bispecific antibodies as derived by structural studies and MD simulations. upper panel: the bivalent14-crs-06 cross-links two spike trimers while the tetravalent 14-H-06 (lower panel) can crosslink up to four spikes. For 14-H-06, two scFvs from CoV2-06 are positioned at one end of the molecule, while two Fab units of CoV2-14 are located at the other end. **e**, Summary of cross-linking potentials by the antibodies reported in this work. *14-H-06 could crosslink three spikes if it binds to three RBDs in the down conformation and one RBD in the up conformation. Otherwise, it could crosslink four spikes.

### Broader coverage of variants by 14-H-06 than the cocktail

We previously identified neutralization-resistant mutation K444R for CoV2-06 and E484A for CoV2-14 and generated SARS-CoV-2 viruses that contain K444R or E484A mutations^9,23^. The K444R virus escaped from CoV2-06 but was neutralized by CoV2-14; the E484A virus escaped from CoV2-14 but was neutralized by CoV2-06 (**Extended Data Fig. 3a-b**). While the two bsAbs and the cocktail (CoV2-06+CoV2-14) neutralized both escaping viruses, their potencies were significantly different. The NT_50s_ for 14-crs-06 against the K444R and E484A viruses were 2.29 μg/ml and 0.83 μg/ml, respectively, which were slightly less potent compared with the NT_50S_ of the cocktail against the K444R (1.02 μg/ml) and E484A (0.59 μg/ml) viruses. In contrast, 14-H-06 neutralized the K444R and E484A viruses with NT_50s_ of 0.23 μg/ml and 0.096 μg/ml, which were 4.4-fold and 6.1-fold more potent compared with the cocktail (**Fig. 4a-b**). Consistent with the neutralization results, the bsAbs bound to the K444R and E484A mutant RBD proteins, while CoV2-06 and CoV2-14 bound to E484A and K444R mutant RBDs, respectively (**Extended Data Fig. 3c**).

**Fig. 4.**
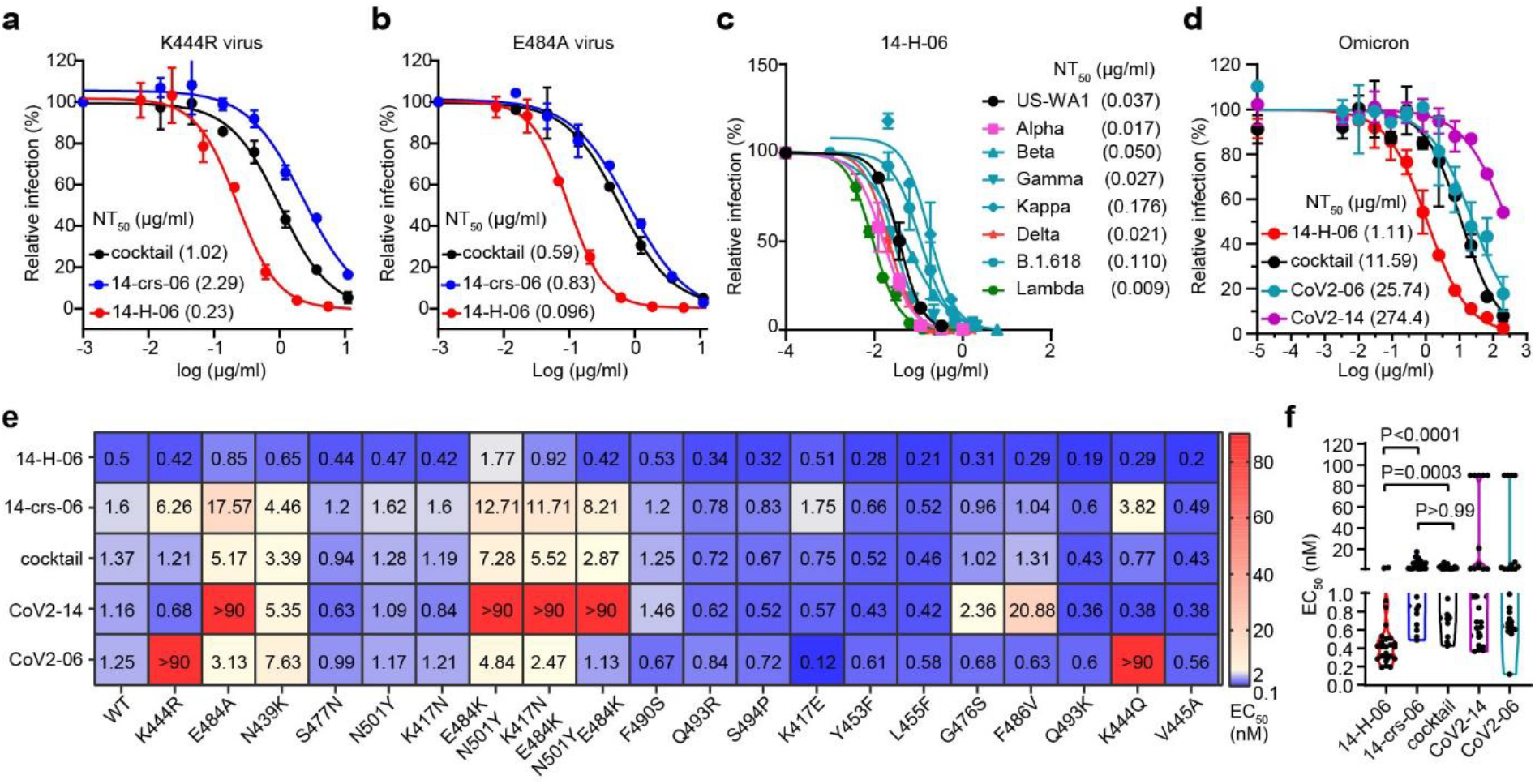
Broad coverage of variants by 14-H-06. **a-b**, Neutralizations of the CoV2-06-resistant SARS-CoV-2 virus with K444R mutation (**a**) and CoV2-14-resistant SARS-CoV-2 virus with E484A mutation (**b**) by indicated bispecific antibodies and the antibody cocktail of CoV2-06 and CoV2-14. The assay is based on the mNeonGreen reporter virus and the NT_50_ values are labeled. **c**, Plaque reduction neutralization test (PRNT) of 14-H-06 against the SARS-CoV-2 US-WA1 strain and recombinant SARS-CoV-2 viruses with the spike replaced by those from indicated variants. The values of neutralizing titers (NT_50_) are labeled. **d**, Neutralizations of the Omicron variant by indicated bispecific antibodies, the antibody cocktail of CoV2-06 and CoV2-14 and two parental antibodies. The assay is based on the mNeonGreen reporter virus and the NT_50_ values are labeled. **e**, Summary of the ELISA binding EC_50_ values of indicated antibodies to the wild type RBD and a panel of 20 RBD mutants. **f**, The violin plot of the EC_50_ values in d. The Kruskal-Wallis test was used for statistical analysis.

We focused on 14-H-06 and evaluated its neutralizing activities against seven SARS-CoV-2 variants using the plaque reduction neutralization test (PRNT) or fluorescent focus reduction test (FFRNT). The complete spike gene from Alpha (B.1.1.7), Beta (B.1.351), Gamma (P.1), Kappa (B.1.617.1), Delta (B.1.617.2), Lambda (C.37), B.1.618 or the Omicron (B.1.1.529) variant was engineered into the backbone of an early clinical isolate USA-WA1/2020 (**Table S2**)^25–27^. Four of these seven variants, including the Beta, Gamma, Kappa, and the B.1.618 variants, carry the E484K or E484Q mutation and were resistant to neutralization by CoV2-14 (**Extended Data Fig. 3d**). Notably, 14-H-06 potently neutralized all the tested variants with the NT_50s_ between 0.009 μg/ml and 0.176 μg/ml, which were in a close range compared with the NT_50_ (0.037 μg/ml) against the US-WA1 strain (**Fig. 4c**). The Omicron (B.1.1.529) variant has 11 RBD mutations, including a G_4_46S mutation within the CoV2-06 epitope and an E484A mutation within the CoV2-14 epitope. The two individual parental antibodies dramatically or almost completely lost neutralizing activity against the Omicron, while remarkably, 14-H-06 neutralized the Omicron with an NT_50_ of 1.11 μg/ml, which is more than 10-fold more potent than the cocktail (**Fig. 4d**).

We further used a collection of 20 mutant RBD proteins to compare the epitope coverages by the two bsAbs, the cocktail, and individual parental antibodies through ELISA titrations (**Extended Data Fig. 4**). These RBDs contain mutations in naturally emerging variants or mutations in escaping viruses selected from two FDA approved antibodies: REGN-10987 and REGN-1093323. Selected RBD mutations reduced the binding activities of individual parental antibodies, such as K444R and K444Q mutations for CoV2-06 and E484A, E484K, and F486V mutations for CoV2-14. Expectedly, the cocktail and the two bsAbs had good coverages of these RBD variants (**Fig. 4e**). Across all the mutants, 14-H-06, but not 14-crs-06, exhibited significantly higher binding activities than the cocktail (**Fig. 4f**), indicating that the IgG-(scFv)_2_ design, but not the CrossMAb design, provides additional advantage for binding to RBD mutants over the cocktail. Together, these data demonstrate that engineering an antibody cocktail into a bsAb using the IgG-(scFv)_2_ design increases the neutralizing potency against SARS-CoV-2 variants and broadens the epitope coverages of RBD mutants.

### *In vivo* protection by 14-H-06

We focused on 14-H-06 to evaluate the *in vivo* efficacy against SARS-CoV-2 and its variants. First, we performed dose range evaluations of 14-H-06 in the Balb/c mice infection model by the CMA4 strain, a mouse-adapted SARS-CoV-2 containing the spike N501Y mutation which represented the Alpha variant^23^ (**Fig. 5a**). Three dose levels (2.5, 0.83 and 0.27 mg/kg) for prophylactic treatment and two dose levels for therapeutic treatment (2.5 and 0.83 mg/kg) were tested. For prophylactic treatment, 14-H-06 reduced the viral loads in the lungs to undetectable levels in 100% (10/10) and 40% (4/10) of mice in the 2.5 and 0.83 mg/kg groups, respectively. Even with the 0.27 mg/kg dose, the geometric mean viral load (4.79-log) was 8.2-fold lower than that from the isotype control group (5.70-log). For 14-H-06 therapy, the geometric mean viral loads (excluding the mice with undetectable viruses) were reduced by 72,766- and 669-fold in the 2.5 and 0.83 mg/kg groups, respectively (**Fig. 5b**). These data demonstrate that 14-H-06 is highly effective for prophylactic and therapeutic treatment against SARS-CoV-2.

**Fig. 5.**
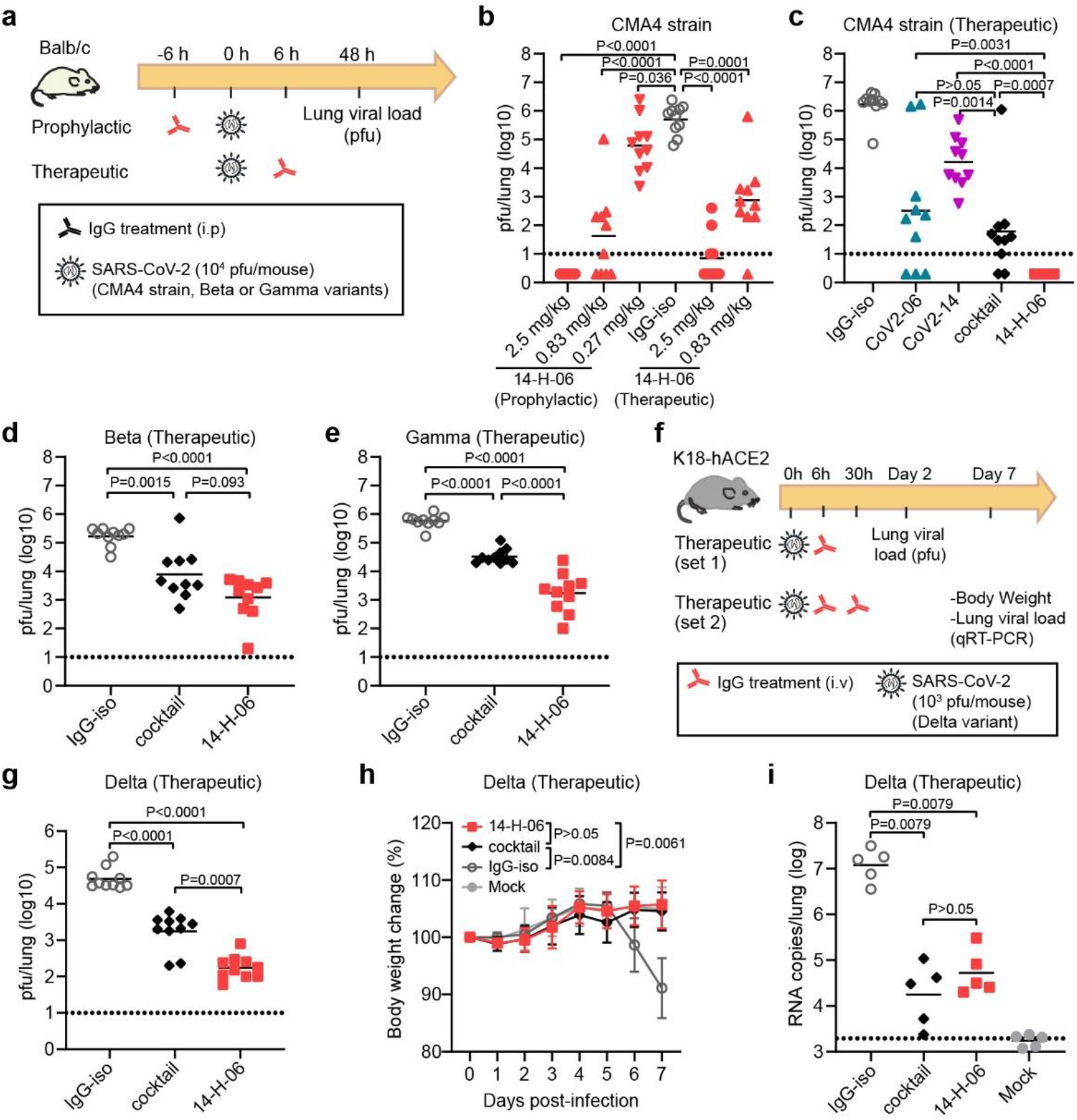
*In vivo* comparisons of 14-H-06 and cocktail against SARS-CoV-2 and variants. **a**, Experimental design for evaluations of the prophylactic and therapeutic effects of antibodies in the Balb/c mouse model of infections. Three SARS-CoV-2 viruses were tested, including a mouse-adapted CMA4 strain containing the spike N501Y mutation and representing the Alpha variant, and the Beta and Gamma variants. n=10 mice for each group. **b**, The viral loads were determined by the pfu assay in the dose-range evaluations of the prophylactic and therapeutic effects of 14-H-06 against the CMA4 virus. **c**-**e**, The viral loads were determined by the pfu assay in the evaluations of the therapeutic effects of indicated antibodies at the dose of 5 mg/kg against the CMA4 virus (**c**), the Beta variant (**d**), and the Gamma variant (**e**). **f**, Experimental design for evaluating the therapeutic effect of 14-H-06 against the Delta variant in the transgenic K18-hACE2 mouse model. In experiment set 1, mice were treated with one dose of antibodies at 6h after infection, and viral loads were measured at 2 days after infection by the pfu assay. In experiment set 2, mice were treated with two doses of antibodies as indicated. The body weight was monitored daily, and the viral loads were measured at 7 days after infection by qRT-PCRR assay. n=10 and 5 mice in each group for set 1 and set 2, respectively. **g**, the lung viral pfu loads for experimental set 1 in panel f. **h-i**, The body weight change (**h**), and the viral RNA load (**i**) for experiment set 2 in panel f. In panels b-e, g, and i, the solid line indicates each group’s geometric mean viral load, and the dotted line indicates the limit of detection (LOD). For statistical analysis, the Mann-Whitney test was used in panels b-e, g, and i; the two-way ANOVA with Tukey’s multiple comparisons was used in panel h.

Next, we compared the therapeutic effects of 14-H-06, the cocktail, and individual parental antibodies against the CMA4 strain. The geometric mean viral load for the cocktail group was 1.79-log, significantly lower than was in the CoV2-14 group (4.21-log) and slightly lower than was in the CoV2-06 group (2.51-log). In contrast, 14-H-06 showed substantially better efficacy than the cocktail and individual parental antibodies, reducing viral loads to undetectable levels for all mice (10/10) (**Fig. 5c**). We also compared the therapeutic effects of 14-H-06 and the cocktail against the Beta and Gamma variants in the Balb/c mouse model following the same experimental design in Fig. 5a. Antibody 14-H-06 significantly reduced the geometric mean lung viral loads by 136-fold for the Beta variant and 333-fold for the Gamma variant compared with the isotype group. A slightly better efficacy against the Beta variant and a more substantial better efficacy for 14-H-06 over the cocktail was observed against the Beta and Gamma variants, respectively (**Fig. 5d-e**). We further compared the therapeutic efficacy of 14-H-06 and the cocktail against the Delta variant in the human ACE2 transgenic mouse (K18-hACE2) model. We performed two sets of experiments to evaluate the therapeutic effects on viral replication (experiment set 1) and mice body weight change (experiment set 2) (**Fig.5f**). In experiment set 1, we treated the mice with one dose of antibodies at 6 h after infection with the Delta variant, and the pfu assay measured viral loads in the lungs. Compared with the isotype group, 14-H-06 reduced the viral load by 278-fold, significantly more potent than the cocktail treatment, which reduced the viral load by 27.8-fold (**Fig.5g**). In experiment set 2, we treated mice at 6 h and 30 h after infection and monitored daily bodyweight. The viral loads were measured seven days post-infection (dpi) by qRT-PCR. The isotype-treated mice showed disease at day 7 post-infection, with an average of 14% body weight loss (**Fig. 5h**) and the geometric mean viral RNA load of 7.2-log (**Fig. 5i**). Treatment with 14-H-06 and the cocktail significantly protected the mice from weight loss (**Fig. 5h**) and reduced the viral RNA loads in the lung (**Fig. 5i**). No significant difference in the lung viral RNA load was observed between 14-H-06 and the cocktail at day 7 (**Fig. 5i**).

Neutralizing antibody levels predict the protection from SARS-CoV-2 infection^28^. We performed a single dose (10mg/kg) pharmacokinetics study in mice to compare the half-life of 14-H-06 with parental antibodies. The half-life for 14-H-06 was 29.2 h compared to 137.4 h and 74.72 h for CoV2-06 and CoV2-14, respectively (**Extended Data Fig. 5a-b**). Thus, the difference in the half-life may complicate the comparison of therapeutic efficacy, particularly in experiment set2 for the Delta variant in the K18-hACE2 model. Taken together, these results demonstrate that 14-H-06 is superior or equivalent to the cocktail for therapeutic treatment of the original SARS-CoV-2 and subsequently emerged Beta, Gamma, and Delta variants in mice.

## Discussion

Neutralizing antibody-based therapies are successful for treating viral infections, yet cocktails are often required to reduce resistance. We have shown that a SARS-CoV-2 bsAb offers advantages in neutralizing activities and spectrum against SARS-CoV-2 variants over the cocktail. Unlike bsAbs using the CrossMAb design, such as CoV-X220 and 14-crs-06, which do not or only slightly increase the neutralizing potency compared to parental antibodies or the cocktail, 14-H-06 significantly increases the neutralizing activity *in vitro* and therapeutic efficacy *in vivo* against SARS-CoV-2 and broadens the coverage of RBD variants. The IgG-(scFv)_2_ design outcompetes the CrossMAb design unlikely via stronger blockage of RBD binding to ACE2, but rather via mechanisms including avidity binding and inter-spike crosslinking. In support of our results, previous studies have shown that multivalent antibodies have greater and broader neutralizing activity than bivalent IgG antibodies^23,29^. Similarly, a SARS-CoV-2 tetravalent bsAb (CV1206_521_GS) uses the DVD-Ig design to combine the RBD- and the NTD-specific antibodies, have demonstrated *in vitro* neutralizing activity that is 100-fold more potent than the cocktail. This DVD-Ig-based bsAb showed good neutralizing coverages of several RBD mutations from some variants^30^; however, its neutralization potency was compromised by the NTD mutations from the Beta and Gamma variants. Nevertheless, rationally designed bsAbs with suitable formats and distinct epitope specificities represent a promising alternative to antibody cocktails for developing COVID-19 therapeutic antibodies.

We directly compared the therapeutic efficacy of 14-H-06 and the cocktail against the spike N501Y mutation-containing CMA4 strain, the Beta, Gamma, and Delta variants *in vivo*. Across all these tested viruses, 14-H-06 has better efficacy than the cocktail regimen. These results support bsAbs as a promising alternative to cocktails for COVID-19 treatment. Although the NT_50s_ of 14-H-06 against the US-WA1 strain, the Alpha, Beta and Gamma variants are in close range (less than 3-fold), 14-H-06 performed better against the CMA4 strain than the Beta and Gamma variants *in vivo*. Notably, antibody Fc-mediated effector functions are required for optimal therapeutic protections against SARS-CoV-2 in mice^31^. Antibody 14-H-06 is engineered using the IgG-(scFv)_2_ design, which is an effector function-competent format^32^. However, it is possible that the effector functions of 14-H-06 have been compromised against the Beta and Gamma variants as a result of the reduced binding for the two Fab14 units to the E484K mutation-containing spikes. The two scFv06 units of 14-H-06 resist the E484K and other mutations in the spike proteins of Beta and Gamma variants. Yet, it is unclear whether the effector functions can be supported in this model of binding. Therefore, choosing an antibody less affected by viral mutations as the IgG backbone for engineering the IgG-(scFv)_2_ format of bsAbs may mitigate the risk of losing Fc-mediated effector functions. Systematic investigation on whether and how bsAb designs affect the Fc-mediated effector functions will provide further insights to guide the development of bsAb-based therapeutic antibodies against SARS-CoV-2.

The IgG-(scFv)_2_ design is a promising platform and has been used for developing more than ten-clinical stage bsAb candidates^33^. Antibody 14-H-06 expresses in high yield (>0.5 g/L) in transient expression and assembles homogenously, suggesting suitable early developmentability profiles. It is noted that 14-H-06 has a shorter half-life than the parental antibodies in mice, which may have limited its therapeutic advantage over the cocktail *in vivo*. The half-life of 14-H-06 may be extended by introducing the M252Y/S254T/T256E (YTE) mutations into the Fc region^34^, or by optimizing the antibody sequence toward favorable physical and chemical properties^35^; and the improved half-life could maximize the therapeutic potential of 14-H-06. The enhanced efficacy of 14-H-6 over the cocktail demonstrated in the *in vitro* and *in vivo* comparisons clearly support the potential to extend the application of the IgG-(scFv)_2_ design to other SARS-CoV-2 antibody cocktails.

In summary, we have engineered two formats of bsAbs and compared them with parental antibodies and the cocktail in a panel of *in vitro* and *in vivo* assays. Our results demonstrate the advantages of a bsAb design over the cocktail in neutralization potency and spectrum. This proof-of-concept study supports that the bsAb approach and the IgG-(scFv)_2_ design can be adapted to broader applications in the development of cost-effective and more efficacious antibody therapies on the basis of antibody cocktails for treating viral infections including SARS-CoV-2.

## Acknowledgments

We thank Dr. Georgina Salazar for her editorial support and Dr Chong Wai Liew for advice. The MD simulations were performed on ASPIRE-1 of the National Supercomputing Centre, Singapore (https://www.nscc.sg).

## Author contributions

Z.K., P.G., and Z.A. conceived the study. Z.K. identified the neutralizing antibody cocktail. P.G., Z.K. and H.S. engineered and produced the bispecific antibodies. X.X, Y.L., A.E.M., J.Z. and V.D.M. performed neutralization and mouse studies. J.L., A.E.S. and B.C.G. performed structural studies. Z.K. and X.Y. generated the RBD mutant proteins. X.L. and X.F. performed mouse PK study. W.X. prepared the Fab. H.D., and H.B. provided support with cell culture and transfection. N.Z., J.L., P-Y.S. and Z.A. supervised the study. Z.K. and P.G. wrote the manuscript with input from the team. X.X., J.L., X.Y., N.Z., J.L., P-Y.S. and Z.A. reviewed and edited the manuscript.

## Data availability

Data associated with figures are available from the corresponding authors upon reasonable request. Structures and structure factors reported in this work have been deposited with the PDB with accession codes 7WPH (Fab-06-RBD complex) and 7WPV (Fab-14). Source data are provided with this paper.

## Material and methods

### Cells, virus and proteins

Expi293F™ cells (GIBCO, cat#100044202) were maintained in Expi293™ Expression Medium without fetal bovine serum (FBS). Vero (ATCC^®^ CCL-81) and Vero E6 cells (ATCC, CRL-1586) were maintained in Dulbecco’s modified Eagle medium (DMEM) supplemented with 10% FBS. The wild-type and K444R and E484A mutations of mNeonGreen SARS-CoV-2 viruses were generated as previously described^36^. The chimeric SARS-CoV-2 viruses with spike gene replaced with B1.1.7, P.1, and B.1.351 linage spike gene were described previously^23^. The chimeric SARS-CoV-2 viruses with spike gene replaced with B.1.617.1, B.1.617.2, B.1.617.2-2, B.1.618 and the Omicron (B.1.1.529) linage spike gene were prepared from clinical strain USA-WA1^36^. Summary of spike mutations of the variants were listed in **Table S1**.The biotinylated SARS-CoV-2 S protein was purchased from Acro Biosystem (Cat# SPN-C82E9-25ug). The His-tagged RBD (RBD-His) protein of SARS-CoV-2 was purchased from Sino Biological (Cat: 40592-V08B). The Fc-tagged wild-type and mutant RBDs mentioned were generated as described previously^23^. The RBD for crystallography harbours a 8xHis tag and is fused to a Maltose Binding Protein via a TEV protease cleavage sequence and was produced from Expi293™ cells. Protein purification was carried out in three steps: an IMAC purification using a HisTrap Ni-NTA column (Cytiva) followed by a TEV cleavage. A reverse IMAC purification was conducted to separate the MBP moiety from the soluble free RBD. The RBD was further purified by size-exclusion chromatography using a S200 16/60 column (Cytiva) pre-equilibrated in phosphate buffered saline at pH 7.2.

### Engineering and production of bsAbs

Plasmids encoding heavy chain, light chain and scFv of CoV2-06 and CoV2-14 were constructed and described previously^9^. For engineering 14-H-06, a similar approach was used as described in a previous study^37^. Briefly, the scFv of CoV2-06 was fused to the C-terminus of CoV2-14 heavy chain with a (G_4_S)_3_ linker to generate 14-H-06 heavy chain plasmid. The bsAb 14-H-06 was expressed by co-transfection of the modified heavy chain and the CoV2-14 light chain plasmids into Expi293F cells. For engineering of 14-crs-06, the CrossMab^CH1-CL^ construct was used as described previously^38^. On one arm, the S354C and T366W mutations were introduced into the heavy chain CH3 region of CoV-06 to generate the hole. This modified heavy chain was paired with the CoV2-06 light chain. On the other arm, the mutations Y349C, T366S, L368A and Y407V mutations were introduced into the heavy chain CH3 region of CoV-14 with the crossover between the CH1 domain and the CL domain of the light chain of CoV2-14. The 14-crs-06 antibody was expressed by co-transfection of four plasmids into Expi293F cells. After 7 days of culture, antibodies were purified using the Protein A resin (Repligen, CA-PRI-0100). All the antibody preparations were reconstituted in phosphate-buffered saline (PBS) buffer for the studies. For the SEC assay, purified antibodies were analyzed on the ÄKTA pure system with the Superpose 6 increase 10/300GL column in PBS buffer. About 100 μg of each antibody was used for each loading. The UNICORN 7.0 software was used to data analysis and exporting.

### In-tandem BLI binding assays

An in-tandem BLI-based binding assay was performed on the Pall ForteBio Octet RED96 system. The RBD-His (1 μg/ml) was loaded onto the Ni-NTA biosensors for 300 seconds. The loaded biosensors were dipped into the first antibody solutions (400 nM) for 300 seconds for the formation of the antibody-antigen complex. The sensors were then dipped into the second antibody solutions (100 nM) for 300 seconds for competition binding. ForteBio’s data analysis software was used to export data, and the binding profile was processed by GraphPad prism 8 Software.

### Antibody affinity and avidity assays

Kinetic analysis was performed using a Pall ForteBio Octet RED96 system. For the affinity assays, antibodies were used as ligands to and loaded onto the Protein A biosensors, at 2 μg/ml for 300s. Following 10s of baseline in kinetics buffer, the loaded biosensors were dipped into serially diluted (0.14–100 nM) RBD-His protein for 300 seconds for association. The sensors were then dipped into a kinetic buffer for 600 seconds to record dissociation. Kinetic buffer without antigen was set to correct the background. For the avidity assays, RBD-His was as ligand and loaded onto the Ni-NTA biosensors at various concentrations (40, 200 and 1000 ng/ml) for 300s. Following 10s of baseline in kinetics buffer, the loaded biosensors were dipped into serially diluted (0.14–100 nM) antibodies 300s for association, then dipped into kinetics buffer 400s for dissociation. ForteBio’s data analysis software was used to fit the K_D_ data using the global fitting method.

### The BLI sandwich assay for testing multivalent binding to RBD

The purified antibodies were tested for their abilities to simultaneously binding to multiple RBD domains on the Octet RED96 system. The RBD-His (5 μg/ml) was captured on the Ni-NTA biosensors for 300 seconds. After capture, the biosensors were dipped into antibody solutions (200 nM) for 300 seconds, and finally to the RBD-Fc solution (200 nM) or PBS control for 300 seconds. The binding responses were recorded for all incubation steps. Last step association (dissociation) was calculated by subtraction of PBS signal from the RBD-Fc binding.

### Crystallization

The Fab-06 and RBD proteins were mixed in a 1:1.2 molar ratio and incubated on ice for 10 minutes, followed by size-exclusion chromatography using a S200 16/60 column (Cytiva) in PBS. The complex peak was pooled and concentrated to 11 mg/ml for crystallization assays which were set up with commercial screening kits (JCSG-plus™ & Morpheus® from Molecular Dimensions; Index™ & PEG/Ion Screen™ from Hampton Research) using a mosquito crystallization robot (TTP Labtech). A thin plate-shaped crystal was obtained in JCSG-plus™ condition A5 (0.2 M magnesium formate dihydrate, 20% w/v PEG 3350) with a protein to buffer ratio of 2 : 1 after 13 days. The crystal was subsequently fished with a nylon loop and flash-frozen in liquid nitrogen and shipped to synchrotron for remote data collection (MXII, ANSTO’s Australian Synchrotron). X-ray diffraction images were integrated and scaled using XDS^39^. Molecular replacement was done via Phaser^40^ using Fv, CH_1_/C_L_, and RBD from PDB accession code 7C01 as three search components. Structure refinement was performed using both Buster^41^ and Phenix Refine^42^ interspersed with manual model correction using Coot^43^. Complex between Fab14 and RBD proteins were also prepared and set up for crystallization in the same manner. Crystals were obtained in 0.1 M Lithium Chloride, 30% (w/v) PEG 4000 with a protein to buffer ratio of 2 : 1 after 7 days. However only Fab14 was present in the crystal. Data collection and refinement statistics for the Fab06-RBD complex and free Fab14 crystal structure are presented in Table S1. Both structures were deposited on Protein Data Bank with accession number 7WPH (Fab-06-RBD complex) and 7WPV (Fab-14).

### Antibody blocking of RBD and ACE2 interaction

The Fc-tagged RBD proteins (4 μg/ml) were captured on the protein A biosensor for 300s. Then, the sensors were blocked by a control Fc protein (150 μg/ml) for 200s to occupy the free protein A on the sensor. The serially diluted antibodies (0.041~30 nM) were then incubated with the sensors for 200s to allow antibody and RBD binding. After 10s of baseline in kinetics buffer, the sensors were dipped in to the ACE2 solution (10μg/ml) for 200s to record the response signal. For analysis of the IC_50_ of the blocking, the ACE2 response values were normalized to the starting points. The blocking percentages at each concentrations were calculated as: (normalized ACE2 response of isotype antibody-normalized ACE2 response of tested antibody)/ normalized ACE2 response of isotype antibody *100. The dose-blocking curves were plotted and the blocking IC_50_ values were calculated by nonlinear fit in the GraphPad prism 8 Software.

### Neutralization assays

All SARS-CoV-2 manipulations were conducted at the Biosafety Level-3 facility with the approval from the Institutional Biosafety Committee at the University of Texas Medical Branch. The neutralizing activities of antibodies against SARS-CoV-2 and two escape mutant strains (K444R and E484A) were measured as previously described using mNeonGreen (mNG) reporter viruses^23^. Briefly, 1.2 ×10^4^ Vero cells were plated into each well of a black clear flat-bottom 96-well plate (Greiner Bio-One; Cat# 655090). The cells were incubated overnight at 37 °C with 5% CO_2_. On the following day, serially diluted antibodies were mixed with an equal volume of virus. After 1 h incubation at 37°C, the antibody-virus complexes were inoculated into Vero cell plates with the final MOI of 2. At 20 h post-infection, nuclei were stained by the addition of Hoechst 33342 to a final concentration of 10 μM. Fluorescent images were acquired using a Cytation 7 multimode reader (BioTek). Total cells (in blue) and mNG-positive cells (in green) were counted, and the infection rate was calculated. The relative infection rates were calculated by normalizing the infection rate of each well to that of control wells (no antibody treatment).

The neutralizing activities of antibodies against SARS-CoV-2 variants were measured using the plaque reduction neutralization test^23^. Briefly, antibodies were 3-fold serially diluted and incubated with 100 plaque forming unit (PFU) of USA-WA1/2020 or mutant SARS-CoV-2. After 1 h incubation at 37 °C, the antibody-virus mixtures were inoculated onto a monolayer of Vero E6 cells pre-seeded on 6-well plates on the previous day. After 1 h of infection at 37°C, 2 ml of 2% SeaPlaque™ Agarose (Lonza) in DMEM containing 2% FBS and 1% penicillin/streptomycin (P/S) was added to the cells. After 2 days of incubation, 2 ml of 2% SeaPlaque™ Agarose in DMEM containing 2% FBS, 1% P/S and 0.01% Neutral Red (Sigma) were added on top of the first layer. After another 16 h of incubation at 37°C, plaque numbers were counted. The dilution concentration that suppressed 50% of viral plaques was defined as PRNT_50_.

### Molecular docking and MD simulations

An intial model for the CoV2-14 scFv-RBD complex was obtained using the HADDOCK 2.4 webserver^44^ by providing the experimental Fab-CoV2-14 structure (this work) and the RBD (PDB access code: 7CJF) X-ray structures as input. An atomic model for the tetravalent bsAb 14-H-06 was built by placing one CoV2-06 scFv molecule at each of the C-terminal ends of the CoV2-14 IgG molecule. A (G_4_S)_3_ linker was then added using MODELLER^45^ to connect each of these CoV2-06 scFv to the CH3 domains of the CoV2-14 IgG. The initial model for the complex between 14-H-06 with four RBD molecules (one RBD bound for each of the four paratopes of IgG-scFv bsAb 14-H-06) was subjected to MD simulations using NAMD 2.12^46^. The system was simulated in a water box where the minimal distance between the solute and the box boundary was 20 Å along all three axes. The charges of the solvated system were neutralized with counter-ions, and the ionic strength of the solvent was set to 150 mM NaCl using VMD^47^. The final system contains over 1.2 million atoms, including proteins, water molecules, and ions. It was subjected to conjugate gradient minimization for 10,000 steps, subsequently heated to 310 K in steps of 5 ps. The system was equilibrated for 5 ns with the backbone atoms constrained by a harmonic potential of the form U(x)=k(x-x_ref_)^2^, where k is 1 kcal mol^−1^Å^−2^ and x_ref_ is the initial atom coordinates. The equilibrated system was simulated for 50 ns under the NPT ensemble assuming the CHARMM^36^ force field for the protein^48^ and assuming the TIP3P model for water molecules. Structure analysis and image production were made using PyMOL (https://pymol.org, Schrödinger Inc.) and COOT^49^.

### ELISA binding titrations of antibodies to RBD mutants

The RBD proteins were coated on Corning high binding assay plates with a concentration of 2 μg/ml at 4°C overnight and blocked with 5% skim milk at 37°C for 2 hours. Serially diluted antibodies were added at a volume of 100 μl per well for incubation at 37°C for 2h. The anti-human IgG Fab2 HRP-conjugated antibody was diluted 1:5000 and added at a volume of 100 μl per well for incubation at 37°C for 1h. The plates were washed 5 times with PBST (0.05% Tween-20) between incubation steps. TMB substrate was added 100μl per well for color development for 3mins and 2 M H_2_SO_4_ was added 50 μl per well to stop the reaction. The OD_450_ was read by a SpectraMax microplate reader. The data points were plotted using GraphPad Prism8, and the EC_50_ values were calculated using a three-parameter nonlinear model.

### Mouse infection models

The animal study was carried out in accordance with the recommendations for care and use of animals by the Office of Laboratory Animal Welfare, National Institutes of Health. The Institutional Animal Care and Use Committee (IACUC) of University of Texas Medical Branch (UTMB) approved the animal studies under protocol 1802011. A previously described mouse infection model was used to evaluate antibody protection. Female BALB/c mice aged 10-12 weeks (n = 10) were infected intranasally (IN) with 10^4^ PFU of mouse-adapted SARS-CoV-2 CMA4 strain^50^ or the Beta and Gamma variants^23^ in 50 μl of PBS. Animals were injected intraperitoneally (i.p.) with antibodies 6 hours before or 6 hours after viral infection. Two days after infection, lung samples of infected mice were harvested and homogenized in 1 ml PBS using the MagNA Lyser (Roche Diagnostics). The homogenates were clarified by centrifugation at 15,000 rpm for 5 min. The supernatants were collected for measuring infectious viruses by plaque assay on Vero E6 cells.

For mouse study with the Delta variant, the 8-10-week-old female K18-hACE2 mice were ordered from The Jackson Laboratory. In experiment set 1, the mice were infected intranasally with 10^3^ PFU of SARS-CoV-2 Delta spike variant (ref: NT162b2-elicited neutralization of B.1.617 and other SARS-CoV-2 variants. Nature 596, 273–275 (2021).) in 50 μl of PBS. Animals were injected intraperitoneally (i.p.) with antibodies 6 hours and 30 hours after viral infection. The body weight of each mouse was monitored daily. Seven days after infection, lung samples of infected mice were harvested and homogenized in 1 ml PBS for qRT-PCR analysis as indicated in a previous study^23^. In experiment set 2, the mice were infected intranasally with 10^4^ PFU of SARS-CoV-2 Delta variant in 50 μl of PBS. Animals were injected intraperitoneally (i.p.) with antibodies 6 hours after viral infection. The body weight of each mouse was monitored daily. Two days after infection, mouse lung samples were harvested and homogenized in 1 ml PBS for plaque assay as described previously^23^.

### The pharmacokinetics of antibodies in mice

Animal experimental protocols were approved by the Animal Welfare Committee at the University of Texas Medical School at Houston. Seven-week-old female BALB/c (Jackson lab, USA) were randomly divided into three groups (5 mice/group) and were injected by i.p with 10 mg/kg of antibody. After injection, mouse blood were collected at 4, 8, 24, 72 h, and day 5, day 7, and day 10. Mouse tail vein was used for blood collection, and up to 0.01 ml of serum was needed for quantification by ELISA. The mouse blood was collected using a sterile scalpel blade, nick the lateral tail vein. Mouse blood (2-3 drops) were collected into Eppendorf tubes. For mouse serum collection, the blood samples were stored at room temperature for 1 hour, and then centrifuged the samples for 30 min at 15,000 rpm at 4°C. The mouse serum was carefully transferred to the new 0.5-ml Eppendorf tubes, and stored them at −20°C until assay. The indirect ELISA was used to quantify serum antibody levels. Briefly, the 96-well plates were coated with the wild type RBD antigen for quantitation of CoV2-06 and CoV2-14 concentrations, and the E481A RBD antigen for quantitation of 14-H-06. Antigens were coated at the concentration of 2 μg/ml in PBS (pH 7.2) and incubated at 4 °C overnight. Plates were blocked with PBS supplemented with 3% BSA at room temperature for 1 h. The mouse sera were diluted 400x for incubation for with plates for 2 h at room temperature. The HRP-conjugated goat anti-human IgG-F(ab’)_2_ was used as the secondary antibody and incubated at room temperature for 1 h. The plate washing, color development steps were the same as described above in ELISA titrations. For analysis of the half-life, the Phoenix 64 WinNonlin (8.3.3.33) software (Certara) was used according to instructions.

## Extended Data Figures

**Extended Data Fig. 1.**
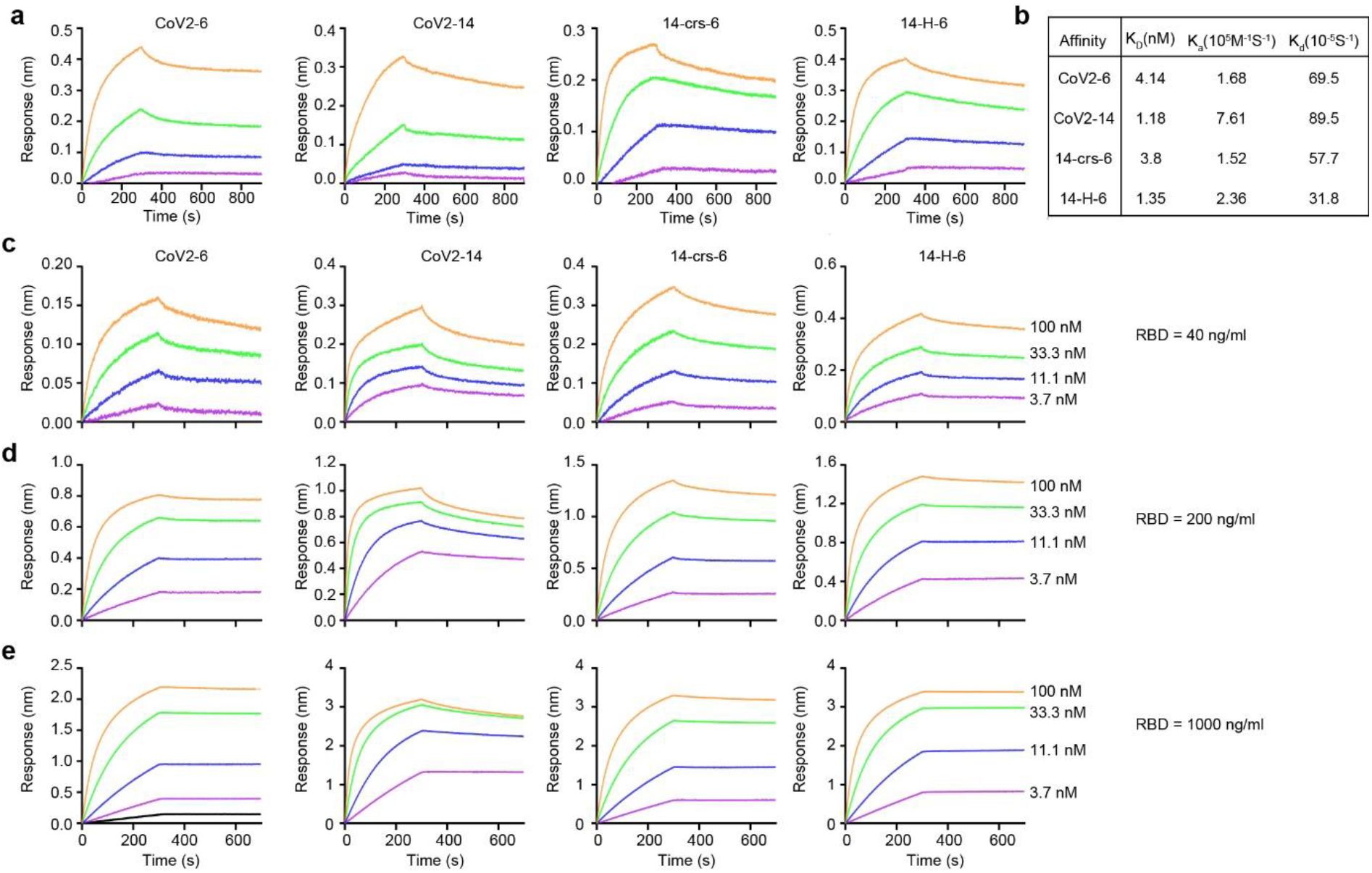
Kinetic bindings of the BLI-based affinity and avidity assays. **a**, The sensorgrams of affinity binding for indicated antibodies. Antibodies were immobilized onto the protein A biosensors and the RBD-His was in solutions. **b**, Summary of the affinity binding (K_D_), the association (K_on_) and the dissociation (K_dis_) parameters. **c**-**e**, The sensorgrams in the avidity binding for indicated antibodies. The RBD-His was immobilized onto the Ni-NTA biosensors at concentrations of 40 ng/ml (**c**), 200 ng/ml (**d**) and 1000 ng/ml (**e**) and indicated antibodies were in solutions.

**Extended Data Fig. 2.**
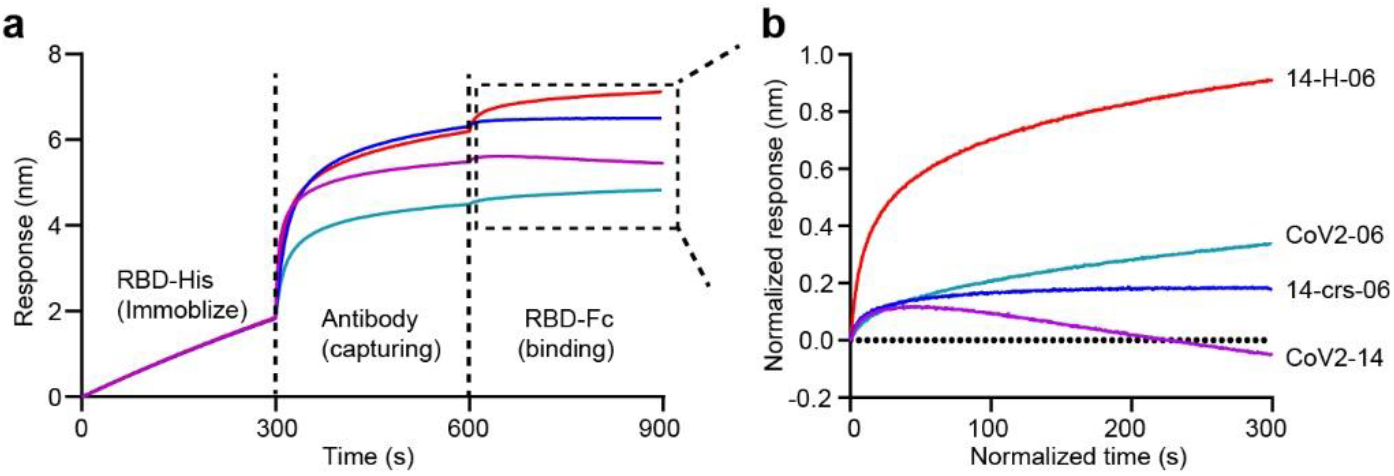
Multivalent binding to RBD by 14-H-06. Simultaneous binding of antibodies to multiple RBDs was determined by a BLI sandwich assay. **a**, The sensorgrams showing the immobilization of RBD-His for 300s, the capturing of indicated antibodies for 300s and the following binding by RBD-Fc for 300s. The binding to RBD-Fc shown in the dashed box was normalized and shown in the panel **b**.

**Extended Data Fig. 3.**
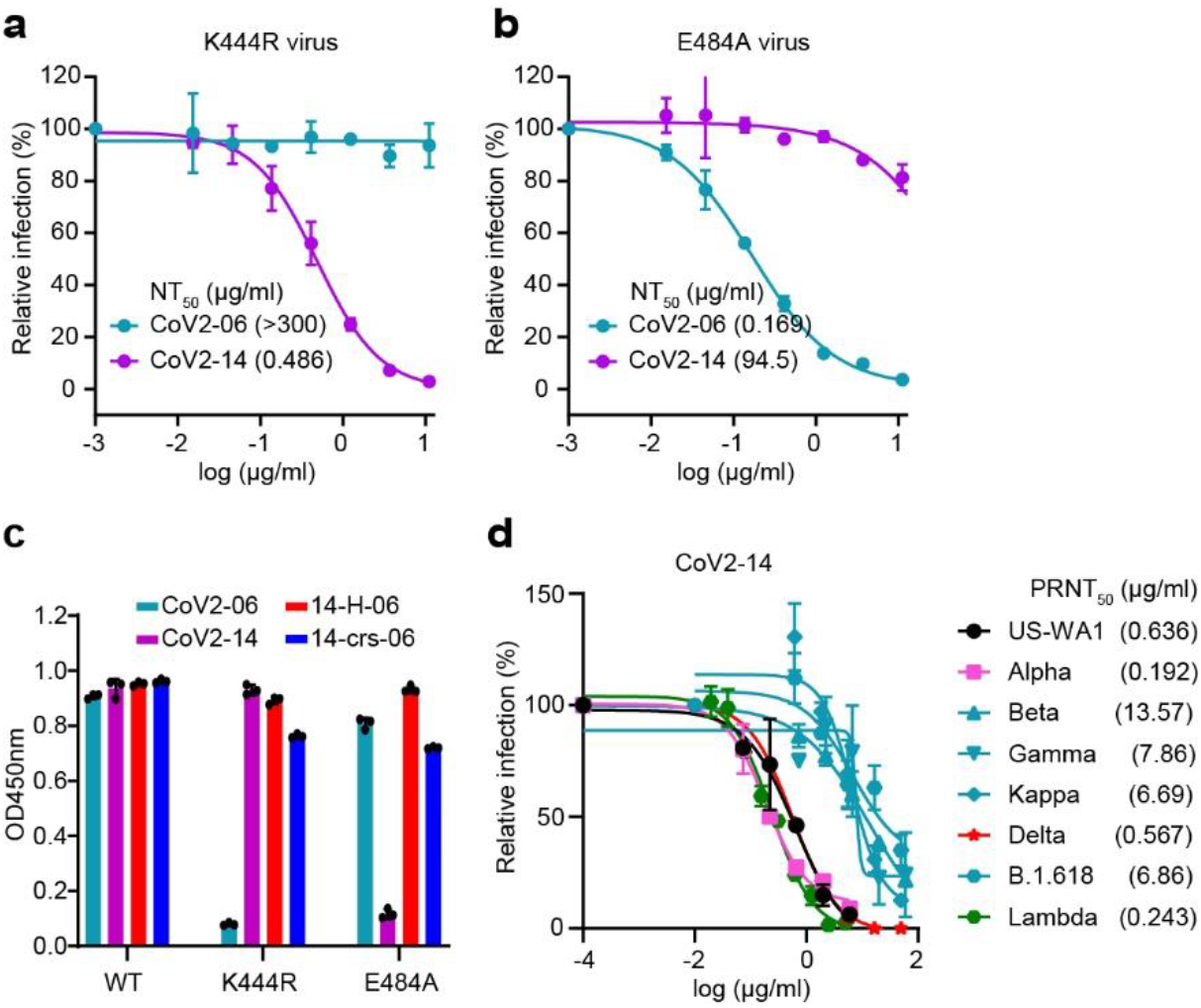
Additional binding and neutralizing characterizations of antibody against the variants. (**a-b**) Neutralizations of SARS-CoV-2 virus with K444R mutation (**a**) and E484A mutation (**b**) by CoV2-06 and CoV2-14. The assay is based on the mNeonGreen reporter virus and the NT_50_ values are labeled. **c**, ELISA binding to the WT RBD and the K444R and E484A RBD mutants by indicated antibodies. **d**, PRNT of CoV2-14 against the SARS-CoV-2 US-WA1 strain and indicated SARS-CoV-2 variants. The PRNT_50_ values are labeled.

**Extended Data Fig. 4.**
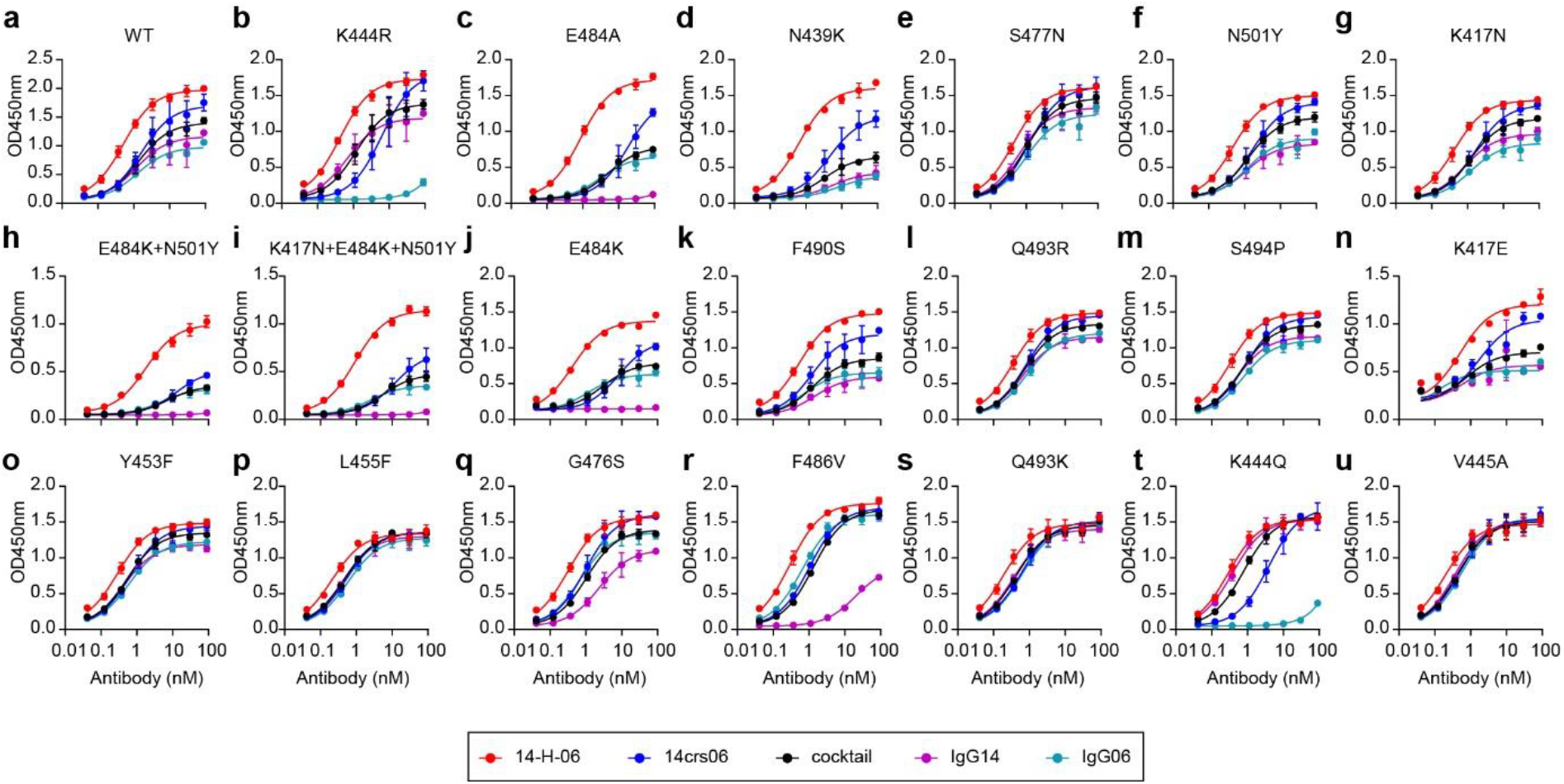
ELISA bindings of bsAbs, individual antibodies and the cocktail to wild type and mutant RBDs. **a**-**u**, ELISA titrations of indicated antibodies to immobilized WT RBD and RBD mutants. Data points are from duplicate wells.

**Extended Data Fig. 5.**
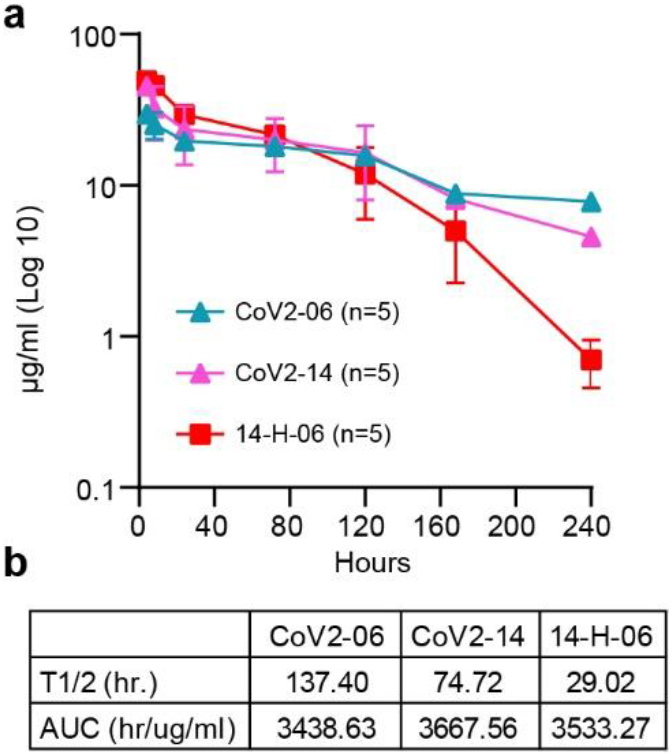
Antibody pharmacokinetics in mice. **a**, The serum concentrations of injected antibodies at multiple time points (4, 8, 24, and 72 hours 5, 7, 10 days) post injection were quantified by ELISA. **b**, Pharmacokinetics parameters were calculated by non-compartmental analysis using Phoe^ni^x 64 WinNonlin (8.3.3.33) software (Certara).

**Table S1.** Data collection and refinement statistics for crystal structures.

|  | **Fab06-RBD Complex** | **Fab14** |
| --- | --- | --- |
| Wavelength (Å) | 0.9464 | 0.9464 |
| Resolution range (Å)<sup>a</sup> | 47.55 - 2.89 (2.993 - 2.89) | 48.03 - 2.46 (2.548 - 2.46) |
| Space group | P 2<sub>1</sub>2<sub>1</sub>2 | C2 |
| Unit cell a,b,c (Å) | 50.17, 266.35, 112.61 | 77.7, 71.22, 96.76 |
| α,β,γ (°) | 90, 90, 90 | 90, 99.07, 90 |
| Total reflections | 363812 (35178) | 130818 (11319) |
| Unique reflections | 34882 (3389) | 18867 (1696) |
| Multiplicity | 10.4 (10.4) | 6.9 (6.7) |
| Completeness (%) | 99.81 (98.86) | 98.83 (89.35) |
| Mean I/sigma(I) | 8.31 (1.08) | 13.60 (1.81) |
| Wilson B-factor | 63.19 | 53.73 |
| R<sub>merge</sub> | 0.3153 (2.504) | 0.1281 (1.076) |
| R<sub>meas</sub> | 0.3316 (2.632) | 0.1385 (1.165) |
| R<sub>pim</sub> | 0.1006 (0.7924) | 0.05213 (0.4418) |
| CC<sub>1/2</sub> | 0.994 (0.48) | 0.997 (0.765) |
| CC* | 0.998 (0.806) | 0.999 (0.931) |
| Reflections used in refinement | 34874 (3388) | 18855 (1694) |
| Reflections used for R-free | 1743 (170) | 943 (85) |
| R<sub>work</sub><sup>b</sup> | 0.2287 (0.3623) | 0.2104 (0.3511) |
| R<sub>free</sub><sup>b</sup> | 0.2694 (0.4140) | 0.2563 (0.3683) |
| CC<sub>(work)</sub> | 0.938 (0.656) | 0.948 (0.795) |
| CC<sub>(free)</sub> | 0.888 (0.374) | 0.907 (0.788) |
| Number of non-hydrogen atoms | 9536 | 3384 |
| protein | 9411 | 3310 |
| NAG | 28 |  |
| water | 97 | 74 |
| Protein residues | 1242 | 438 |
| RMS(bonds) (Å) | 0.012 | 0.010 |
| RMS(angles) (°) | 1.55 | 1.27 |
| Ramachandran favored (%) | 95.43 | 95.39 |
| Ramachandran allowed (%) | 4.57 | 4.61 |
| Ramachandran outliers (%) | 0.00 | 0.00 |
| Rotamer outliers (%) | 0.66 | 0.00 |
| Clashscore | 13.06 | 8.74 |
| Average B-factor | 67.21 | 57.01 |
| protein | 67.34 | 57.00 |
| water | 49.33 | 57.17 |

**Table S2.** Engineered mutations in the spike region of recombinant SARS-CoV-2 variants.

| **Variants** | **Spike mutations** |
| --- | --- |
| Alpha<br>(B.1.1.7) | Δ69-70, Δ145, N501Y, A570D, D614G, P681H, T716I, S982A, and D1118H |
| Beta<br>(B.1.351) | D80A, D215G, Δ242-244, K417N, E484K, N501Y, D614G, and A701V |
| Gamma<br>(P.1) | L18F, T20N, P26S, D138Y, R190S, K417T, E484K, N501Y, D614G, H655Y, T1027I, and V1176F |
| Kappa<br>(B.1.617.1) | G142D, E154K, L452R, E484Q, D614G, P681R, Q1071H, and H1101D |
| Delta<br>(B.1.617.2) | T19R, G142D, L452R, T478K, D614G, P681R, and D950N |
| Lambda | G75V, T76I, Δ246-252, D253N, L452Q, F490S, D614G, and T859N |
| B.1.618 | H49Y, Δ145-146, E484K, and D614G |
| Omicron<br>(B.1.1.529) | A67V, del69-70, T95I, Δ142-144, Y145D, del211, L212I, ins214EPE, G339D, S371L, S373P, S375F, K417N, N440K, G446S, S477N, T478K, E484A, Q493R, G496S, Q498R, N501Y, Y505H, T547K, D614G, H655Y, N679K, P681H, N764K, D796Y, N856K, Q954H, N969K, L981F |

